# Coverage Geometry and Heuristic Discovery Times in Protein Sequence Space

**DOI:** 10.64898/2026.09.01.748471

**Authors:** Haziq Moinudeen, Alp Duygu

**Affiliations:** Elizabeth Garrett Anderson Institute for Women’s Health, University College London; Department of Chemistry, University College London

**Keywords:** protein sequence space, Hamming geometry, neutral networks, functional density, evolutionary search

## Abstract

Functional protein sequence space is often described either by the sparsity of functional sequences or by the connectivity of neutral networks. These quantities characterize different properties: density measures the fraction of all sequences that are functional, whereas connectivity describes relationships among functional sequences. Neither alone determines how much of sequence space lies close to function.

The *coverage function* measures the proportion of sequence space within a prescribed Hamming distance of functionality and therefore provides a direct geometric measure of local accessibility. Here we connect this geometric framework to characteristic discovery times using an explicitly heuristic model of stochastic exploration. The explored region is represented by an effective Hamming ball, and radial displacement is modelled as an outward-biased substitution process. A numerical illustration, calibrated to an empirical human germline mutation rate, shows how a coverage radius can be converted into a timescale. We then extend the framework to parallel search by defining an overlap-adjusted effective number of trajectories. The purpose is not to predict exact evolutionary waiting times, but to separate the geometric distribution of function from the dynamics by which sequence space is explored.

## 1 Introduction

The enormous size of protein sequence space has framed discussions of molecular evolution since Maynard Smith’s formulation of protein space, which emphasized that evolution proceeds through local mutational steps rather than arbitrary jumps between sequences [1]. Subsequent theoretical and computational work showed that viable protein sequences can form extended neutral networks connected by single mutations [2, 3, 4].

Much of this literature concerns the connectivity, robustness, or traversability of viable regions: whether sequences retaining a fold or function can be connected through local mutational steps [3, 4]. Geometry alone, however, cannot determine evolutionary accessibility. A functional network may be highly connected while occupying an exponentially small fraction of sequence space, or it may consist of many disconnected components while nevertheless lying close to most sequences. We therefore consider the *coverage function*: the fraction of total sequence space within a prescribed Hamming distance of the functional set.

Related local-neighbourhood questions have been considered previously. In RNA, Schuster emphasized that common structures can be encountered within comparatively small neighbourhoods of sequence space [5]. In protein models, Aita et al. explicitly examined neutral networks using Hamming distance and the radii of sequence-space balls required to encounter different structural classes [4]. The raw cardinality *q*^*L*^ is itself insufficient to characterize evolutionary exploration; population-level arguments likewise depend strongly on how exploration is defined [6]. Here we abstract away from the internal structure of any particular neutral network and ask for an arrangement-independent relation between global functional density and the fraction of sequence space lying within distance *r* of function.

Having isolated this geometric quantity, we ask how long stochastic exploration takes to reach its characteristic discovery radius. We also consider parallel exploration by related descendants. Shared ancestry makes their trajectories correlated: population size increases the breadth of the search cloud but does not multiply radial displacement along any one lineage. This leads to an overlap-adjusted effective trajectory count rather than a multiplier based on cumulative census alone.

The calculations that follow are intentionally heuristic and represent the exploration cloud by an effective Hamming ball. For the mammalian illustration, we derive the amino-acid-changing mutation rate from an empirical human germline SNV rate and the standard genetic code. Figure 1 summarizes the conceptual progression from functional density and coverage geometry to radial dynamics, discovery time, and effective parallelism.

**Figure 1.**
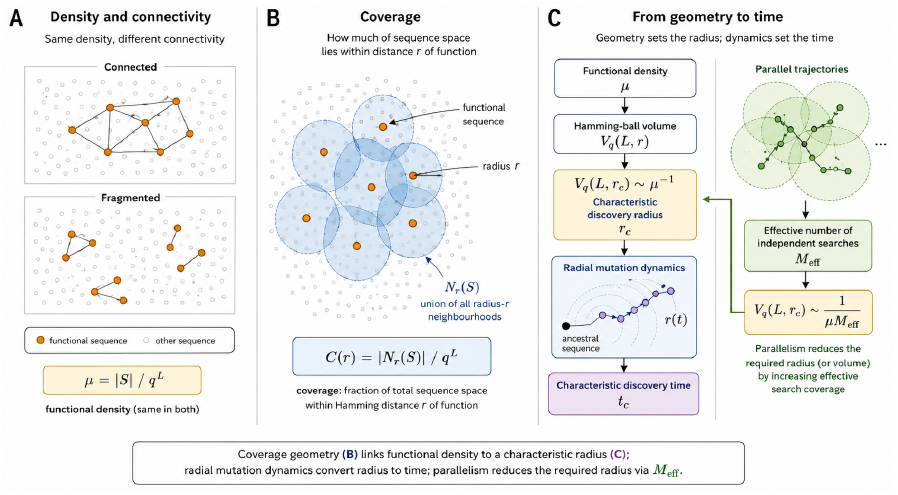
Conceptual framework linking functional density, coverage geometry, and discovery time in protein sequence space. (A) Functional density alone does not determine connectivity: sets containing the same fraction of functional sequences may form a connected network or multiple fragmented components. (B) Coverage *C*(*r*) is the fraction of total sequence space contained in the union of radius-*r* Hamming neighbourhoods around the functional set. (C) The characteristic radius *r*_*c*_ is obtained from the coverage geometry and converted to a characteristic time through the radial mutation dynamics. Effective parallelism reduces the required search volume, and therefore the required radius, through *M*_eff_. The two-dimensional arrangements are schematic only and do not represent literal Euclidean embeddings of the high-dimensional Hamming space. The schematic was generated with assistance from ChatGPT (OpenAI) from an author-specified conceptual design and was reviewed by the authors for mathematical and conceptual accuracy.

## 2 Coverage Geometry

Let 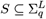 denote the set of functional protein sequences of length *L* over an alphabet of size *q*. The functional density is

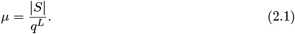

For every nonnegative integer *r*, define the covered neighbourhood

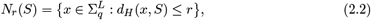

where *d*_*H*_ denotes Hamming distance. The coverage fraction is therefore

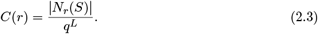

The Hamming ball of radius *r* contains

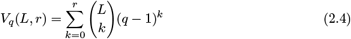

sequences. The covered neighbourhood is a union of at most |*S*| such balls, so the union bound gives

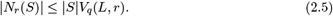

Dividing by the total number of sequences yields the fundamental inequality.

**Fundamental coverage bound**

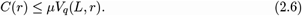

This result is completely independent of the connectivity of the functional network. Suppose now that functional density decreases exponentially with protein length,

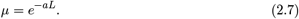

For every fixed radius *r*,

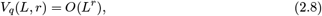

which grows only polynomially with sequence length. Consequently,

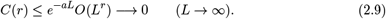

Thus exponentially sparse functionality cannot be compensated by exploration over any bounded mutational radius.

## 3 A Mean-Field Model of Evolutionary Exploration

The previous section concerns only geometry. To relate geometry to evolutionary time we introduce a deliberately simple stochastic model. We assume mutations occur independently at a constant whole-protein mutation rate *U* per generation.

Consider a sequence of length *L* over an alphabet of size *q* that currently lies at Hamming distance *r* from its ancestral sequence. Of its *L* positions, *L − r* retain their ancestral state and *r* contain a non-ancestral state.

A mutation at one of the *L − r* ancestral positions necessarily increases the Hamming distance by one. Therefore,

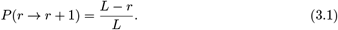

A mutation at one of the *r* non-ancestral positions decreases the Hamming distance only if the new state is the ancestral state. Under uniform substitution among the *q −* 1 states different from the current state,

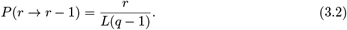

The remaining mutations at non-ancestral positions replace one non-ancestral state with another and therefore leave the Hamming distance unchanged:

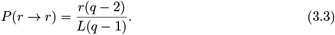

The expected change in Hamming distance per mutation event is consequently

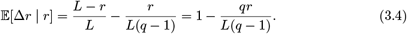

At small Hamming distances, most mutations move the sequence outward, whereas an inward step requires a specific back-substitution at an altered position.

Define

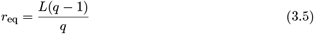

and

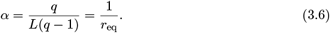

The outward drift vanishes at *r* = *r*_eq_, which is also the expected Hamming distance between two independently sampled random *q*-ary sequences.

Let *R*_*n*_ denote the Hamming distance after *n* mutation events, with *R*_0_ = 0. Taking expectations in Equation (3.4) gives

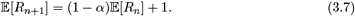

Solving this recurrence yields

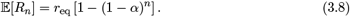

For *n* small relative to *r*_eq_,

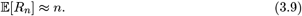

Thus radial displacement is initially approximately linear in the number of mutation events, with the outward drift weakening as the trajectory approaches *r*_eq_. Related work has modelled long-term protein divergence from an ancestral sequence as substitution trajectories through sequence space [7].

The resulting radial dynamics are shown in Figure 2. At the characteristic radius relevant to the illustrative example, the process remains deep in the outward-drift regime. Direct sequence-level Monte Carlo simulation closely follows the analytic expectation over the full trajectory.

**Figure 2.**
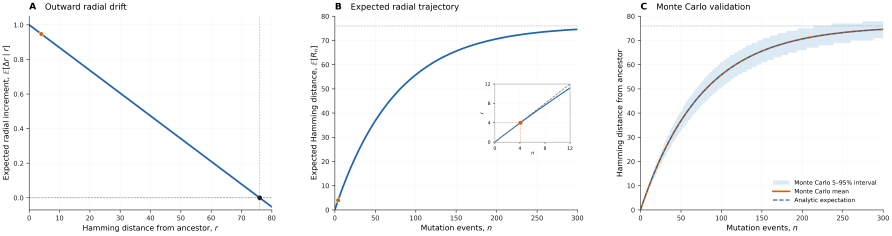
Radial substitution dynamics in *q*-ary protein sequence space for *L* = 80 and *q* = 20. (A) Expected radial increment per mutation event as a function of Hamming distance from the ancestral sequence. The drift is strongly outward at small radius and vanishes at the equilibrium distance *r*_eq_ = 76. The orange point marks the characteristic radius *r*_*c*_ = 4 for the *µ* = 10^*−*11^illustrative example. (B) Analytic expectation of Hamming distance after *n* mutation events, E[*R*_*n*_] = *r*_eq_[1 *−* (1 *−*1*/r*_eq_)^*n*^]. The inset shows the early-time regime, in which radial distance is approximately linear in mutation-event count; *r*_*c*_ = 4 corresponds to *n*_*c*_ *≈* 4.08. (C) Direct sequence-level Monte Carlo simulation of the *q*-state substitution process compared with the analytic expectation. The shaded region denotes the 5th–95th percentile interval across simulated trajectories.

We now make a separate mean-field approximation: the irregular exploration cloud associated with radius *r* is replaced by an effective Hamming ball of volume *V*_*q*_(*L, r*). This does not imply that every sequence in the ball has literally been visited. Assuming functionality is approximately uniformly distributed within this effective region at global density *µ* then gives the discovery criterion below.

## 4 Expected Functional Discoveries

Under the mean-field approximation, the expected number of functional sequences within an effective exploration radius *r* is

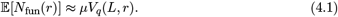

A characteristic discovery radius satisfies E[*N*_fun_(*r*_*c*_)] *≈* 1, or equivalently,

**Characteristic discovery radius**

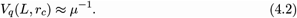

This criterion is purely geometric; mutation rate, generation time, and population dynamics enter only when converting *r*_*c*_ into a timescale.

## 5 From Radius to Time

If mutations occur at rate *U* per generation, then after time *t* the expected number of mutation events along a lineage is

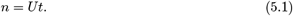

Substituting this into Equation (3.8) gives the expected radial displacement

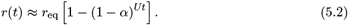

For a characteristic discovery radius *r*_*c*_, inversion of Equation (5.2) gives

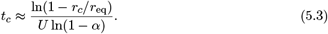

For *r*_*c*_ *≪ r*_eq_, Equation (5.3) reduces to

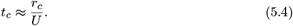

For *r*_*c*_ *≪ r*_eq_, discovery time therefore scales approximately linearly with the required radius.

## 6 Illustrative Estimates

To provide an empirically anchored illustration, consider the random-sequence ATP-binding experiment of Keefe and Szostak [8]. Their selection began from a library of approximately 6 *×* 10^12^proteins containing 80 contiguous randomized amino-acid positions and yielded an estimated frequency of ATP-binding proteins of approximately one in 10^11^random sequences. For the geometric calculation we therefore take *L* = 80, *q* = 20, and *µ* = 10^*−*11^. The value *µ* = 10^*−*11^is phenotype- and assay-specific. It should not be interpreted as a general estimate of the density of all functional proteins in sequence space. Other experimental approaches have produced substantially different estimates for the prevalence of particular folds or activities [9], underscoring that *µ* is an assay- and phenotype-specific parameter rather than a universal density of functional protein.

### Empirical calibration of the amino-acid mutation rate

The dynamical model requires the rate *U* of amino-acid-changing mutations per protein per generation. Kong et al. [10] estimated the human germline single-nucleotide mutation rate as

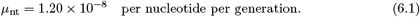

We convert this nucleotide-level rate to the corresponding missense rate using the standard genetic code. The 61 sense codons each admit nine directed single-nucleotide changes, giving 549 possible changes in total. Of these, 134 are synonymous, 392 missense, and 23 nonsense (Table 1).

**Table 1.** Classification of all directed single-nucleotide substitutions from the 61 sense codons of the standard genetic code.

| OUTCOME | NUMBER OF SUBSTITUTIONS | FRACTION |
| --- | --- | --- |
| Synonymous | 134 | 0.2441 |
| **Missense** | **392** | **0.7140** |
| Nonsense | 23 | 0.0419 |
| Total | 549 | 1.0000 |

The resulting missense fraction is

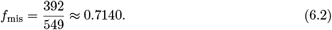

For a protein of length *L*, the amino-acid-changing mutation rate is

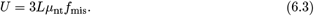

For the *L* = 80 sequence considered below,

**Empirically calibrated whole-protein rate**

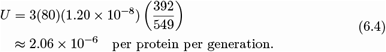

This conversion weights the possible single-nucleotide changes equally; real germline mutation spectra are context dependent. It is therefore a transparent first-order calibration rather than a complete model of coding mutation.

Direct evaluation of Equation (2.4) gives the Hamming-ball volumes shown in Table 2.

**Table 2.** Selected Hamming-ball volumes for *L* = 80 and *q* = 20.

| RADIUS | APPROXIMATE $V_{20}(80, r)$ |
| --- | --- |
| 0 | 1 |
| 1 | $1.52 \times 10^3$ |
| 2 | $1.14 \times 10^6$ |
| 3 | $5.65 \times 10^8$ |
| 4 | $2.07 \times 10^{11}$ |

Since *µ*^*−*1^ = 10^11^,

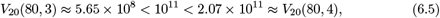

so the discovery criterion is first crossed at integer radius *r*_*c*_ = 4. The radial parameters are

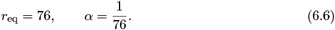

The corresponding number of mutation events is

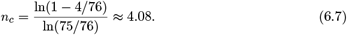

Hence

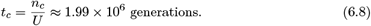

For illustration, these correspond to the calendar times shown in Table 3.

**Table 3.** Illustrative calendar times under two generation-time assumptions.

| ORGANISM | APPROXIMATE TIME |
| --- | --- |
| Bacteria (20-minute generations) | $7.5 \times 10^1$ years |
| Mammal (illustrative 2-year generation time) | $4.0 \times 10^6$ years |

These calculations are intended only as order-of-magnitude illustrations under the stated assumptions. They should not be interpreted as precise biological predictions.

## 7 Parallel Search and Effective Trajectories

Population size does not multiply the rate of radial displacement along a single lineage. It creates parallel trajectories, and only the portions of those trajectories that do not overlap enlarge the explored region. Parallel trajectories need not constitute independent searches: descent from a common ancestor can itself strongly restrict the fraction of protein sequence space explored [11]. A cumulative count of individuals therefore cannot be treated as the same number of independent random initializations.

Let *ℰ*_*i*_(*t*) denote the effective region associated with lineage *i* by time *t*, and let

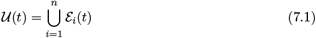

be their combined explored region. We define the overlap-adjusted effective trajectory number by

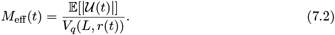

For equal-radius search clouds this quantity obeys

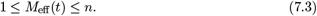

The upper limit describes unrelated, non-overlapping searches; the lower limit describes complete overlap. Shared ancestry and nearby starting points generally place real populations between these limits.

Under the same uniform-density approximation used above, the expected number of functional sequences in the union is

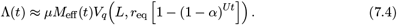

If rare functional encounters are approximated by a Poisson process, then

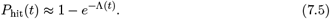

Consequently, the time required to reach a chosen discovery probability *p* is determined implicitly by the following criterion.

**Parallel discovery criterion**

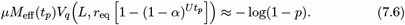

The earlier rule is recovered at *p* = 1 − *e*^*−*1^, for which the right-hand side equals one. If *M*_eff_(*t*) is approximated by a constant *M*, parallelism changes the required volume and hence the critical radius,

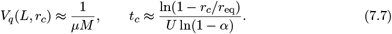

It does *not* justify dividing the single-lineage time directly by *M*.

### Illustrative effect of parallelism

To illustrate the scale of this effect, Table 4 applies the constant-*M* approximation to the Keefe–Szostak example.

**Table 4.** Effect of an illustrative constant effective-trajectory multiplier *M* on the discovery scale. *M* denotes overlap-adjusted effective search multiplicity, not cumulative census size.

| SEARCH ENSEMBLE | TARGET VOLUME | RADIUS | 2-YEAR MAMMALIAN TIME |
| --- | --- | --- | --- |
| Single trajectory | $10^{11}$ | 4 | $4.0 \times 10^6$ years |
| $M = 10^4$ | $10^7$ | 3 | $3.0 \times 10^6$ years |
| $M = 10^{11}$ | 1 | 0 | No exploration required |

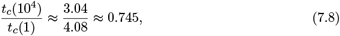

so a 10^4^-fold effective trajectory multiplier reduces the illustrative time by only about 25%, rather than by a factor of 10^4^. At *M* = 10^11^, *µM* = 1, so the expected-one-hit criterion is already met at radius zero in the idealized independent-initialization limit.

The geometric consequences of functional rarity and effective parallelism are summarized in Figure 3. Functional rarity increases the required radius stepwise because Hamming radius is integer-valued, whereas effective parallelism reduces the required radius by increasing the effective search coverage.

**Figure 3.**
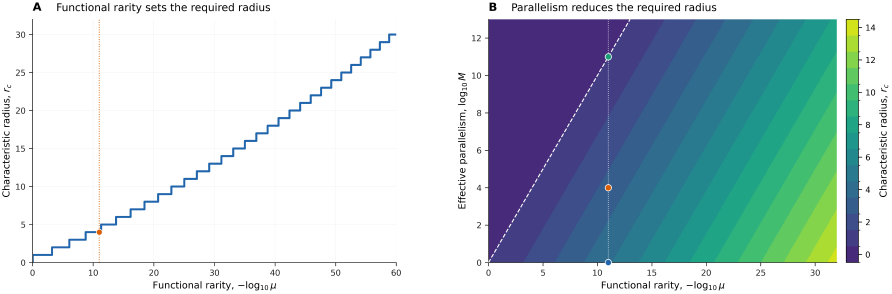
Characteristic discovery radius as a function of functional rarity and effective parallelism for *L* = 80 and *q* = 20. (A) Smallest integer Hamming radius *r*_*c*_ for which *V*_*q*_(*L, r*_*c*_) reaches the order-one mean-field discovery criterion. The staircase structure follows from the discreteness of Hamming radius. The orange marker indicates *µ* = 10^*−*11^, for which *r*_*c*_ = 4. (B) Characteristic radius obtained from *V*_*q*_(*L, r*_*c*_) *≈* 1*/*(*µM*_eff_) as a function of functional rarity and effective parallelism. The vertical dotted line corresponds to *µ* = 10^*−*11^. The three markers show *M*_eff_ = 1, 10^4^, and 10^11^, giving *r*_*c*_ = 4, 3, and 0, respectively. The white dashed diagonal marks *µM*_eff_ = 1; at and beyond this boundary the order-one mean-field criterion requires no radial exploration.

## 8 Discussion

The central result is the separation between geometry and dynamics. Coverage determines the characteristic radius through

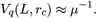

Mutation dynamics then determine the time required to reach that radius.

The mean-field approximation deliberately removes much of the structure known to characterize real genotype–phenotype maps. Functional sequences need not be distributed uniformly: protein structures and functions can occupy highly heterogeneous regions of sequence space, and local mutational tolerance depends strongly on sequence background [12, 13, 14]. This heterogeneity does not affect the arrangement-independent coverage bound, but it limits the interpretation of *µV*_*q*_(*L, r*) as a local discovery expectation.

The radial model is likewise deliberately minimal. Local constraints need not imply permanent confinement: empirical sequence analysis suggests that compensatory substitutions can continually open new directions of protein sequence-space exploration over long evolutionary times [15]. More detailed mutation models can replace the present radial dynamics without altering the coverage criterion.

For populations, parallel exploration is governed by the union of overlapping search regions rather than raw census size. The effective trajectory number *M*_eff_ summarizes this overlap. Increasing *M*_eff_ lowers the volume, and hence radius, required of each trajectory, but does not divide the single-lineage discovery time directly by population size. The restriction imposed by correlated descent provides direct biological motivation for this overlap adjustment [11].

The coverage bound is rigorous, whereas the effective Hamming ball, uniform-density assumption, and temporal calculations are heuristic. Explicit neutral-network models can incorporate correlations, background-dependent viability, and richer substitution dynamics [16]. The present purpose is not to provide precise evolutionary waiting times, but to separate the geometric question of how function is distributed through sequence space from the dynamical question of how that space is explored.

## 9 Computational methods

The *q*-ary Hamming-ball volumes were evaluated directly as

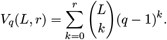

For each parameter combination, *r*_*c*_ was defined computationally as the smallest integer radius satisfying the relevant order-one volume criterion: *V*_*q*_(*L, r*_*c*_) *≥ µ*^*−*1^ for a single effective search and *V*_*q*_(*L, r*_*c*_) *≥* (*µM*_eff_)^*−*1^with effective parallelism. When *µM*_eff_ *≥* 1, *r*_*c*_ = 0. The staircase in Figure 3 therefore reflects the exact integer-valued Hamming-radius calculation rather than interpolation.

The analytic curve in Figure 2 evaluates

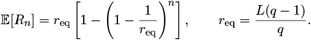

The sequence-level Monte Carlo calculation used *L* = 80, *q* = 20, 20,000 independent trajectories, 300 mutation events per trajectory, and random seed 20260901. Each trajectory began from the same ancestral sequence. At each event, one of the *L* sites was selected uniformly and its current residue was replaced uniformly by one of the other *q* − 1 residue states. Hamming distance from the ancestral sequence was recorded after each event. The plotted Monte Carlo mean is the mean across trajectories, and the shaded interval is the 5th–95th percentile. Numerical calculations and plotting were performed in Python using NumPy and Matplotlib.

## Data and code availability

No new empirical data were generated in this study. All numerical results and data visualizations are generated directly from the equations and algorithms described in the text. The Python code used to calculate *q*-ary Hamming-ball volumes, determine characteristic radii, perform the sequence-level Monte Carlo simulations, and generate Figs. 2 and 3 will be made available with the accompanying source materials.

## Declaration of generative AI and AI-assisted technologies in the manuscript preparation process

During the preparation of this work, the authors used ChatGPT (OpenAI) to assist with manuscript editing, computational code development, and the initial generation of the explanatory schematic in Fig. 1. All mathematical derivations, numerical results, code outputs, figure content, references, and manuscript text were reviewed and edited by the authors, who take full responsibility for the content of the work.

