## Supplementary material for "Coverage Geometry and Heuristic Discovery Times in Protein Sequence Space": figures and code: protein_hamming_geometry.ipynb

```
# %% [code]
#!/usr/bin/env python3
# -*- coding: utf-8 -*-

"""
===============================================================================
V5 — MINIMAL PUBLICATION FIGURES
Coverage Geometry and Heuristic Discovery Times in Protein Sequence Space
===============================================================================

This is a CLEAN, STANDALONE replacement for V4.

Design philosophy
-----------------
The figures contain DATA, not prose.

No:
    - arrow annotations
    - floating parameter labels
    - text boxes
    - labels attached to curves
    - labels attached to points
    - contour labels
    - explanatory sentences inside panels
    - equilibrium labels written over curves
    - decorative callouts

Interpretation belongs in the figure caption.

Outputs
-------
protein_sequence_space_figures_v5/
    Figure_2_radial_substitution_dynamics_v5.png
    Figure_2_radial_substitution_dynamics_v5.pdf

    Figure_3_discovery_geometry_v5.png
    Figure_3_discovery_geometry_v5.pdf

    panels/
        Figure_2A_v5.png/pdf
        Figure_2B_v5.png/pdf
        Figure_2C_v5.png/pdf
        Figure_3A_v5.png/pdf
        Figure_3B_v5.png/pdf

    data/
        hamming_ball_volumes.csv
        monte_carlo_summary.csv
        numerical_summary.txt
"""

# =============================================================================
# IMPORTS
# =============================================================================

from pathlib import Path
from math import comb
import csv

import numpy as np
import matplotlib.pyplot as plt

from matplotlib.colors import BoundaryNorm, ListedColormap
from matplotlib.ticker import MultipleLocator
from mpl_toolkits.axes_grid1.inset_locator import inset_axes


# =============================================================================
# OUTPUT DIRECTORIES
# =============================================================================

ROOT = Path("protein_sequence_space_figures_v5")
PANEL_DIR = ROOT / "panels"
DATA_DIR = ROOT / "data"

for d in [ROOT, PANEL_DIR, DATA_DIR]:
    d.mkdir(parents=True, exist_ok=True)


# =============================================================================
# MODEL PARAMETERS
# =============================================================================

L = 80
q = 20

# Functional density used in manuscript example
MU = 1e-11

# Human germline SNV rate
MU_NT = 1.20e-8

# Exact fraction of directed single-nucleotide changes among sense codons
# that are missense
F_MISSENSE = 392 / 549

# Whole-protein amino-acid-changing mutation rate
U = 3 * L * MU_NT * F_MISSENSE

# Radial equilibrium distance
R_EQ = L * (q - 1) / q

# alpha = 1 / r_eq
ALPHA = 1 / R_EQ


# =============================================================================
# MONTE CARLO PARAMETERS
# =============================================================================

RNG_SEED = 20260901

N_TRAJECTORIES = 20_000
N_EVENTS = 300

LOW_Q = 0.05
HIGH_Q = 0.95


# =============================================================================
# VISUAL CONSTANTS
# =============================================================================

BLUE = "#2166AC"
ORANGE = "#D95F02"
GREEN = "#1B9E77"

BLACK = "#202020"
GREY = "#777777"
LIGHT_GREY = "#D9D9D9"
PALE_BLUE = "#CFE4F1"
WHITE = "#FFFFFF"


# =============================================================================
# MATPLOTLIB CONFIGURATION
# =============================================================================

plt.rcParams.update({

    "font.family": "sans-serif",
    "font.sans-serif": [
        "Arial",
        "Helvetica",
        "DejaVu Sans"
    ],

    "mathtext.fontset": "dejavusans",

    "font.size": 9.5,

    "axes.labelsize": 10.5,
    "axes.titlesize": 11.5,

    "xtick.labelsize": 9,
    "ytick.labelsize": 9,

    "legend.fontsize": 8.2,

    "axes.linewidth": 0.8,

    "axes.spines.top": False,
    "axes.spines.right": False,

    "xtick.major.width": 0.8,
    "ytick.major.width": 0.8,

    "xtick.major.size": 4,
    "ytick.major.size": 4,

    "figure.facecolor": "white",
    "axes.facecolor": "white",
    "savefig.facecolor": "white",

    # Preserve editable text in PDF
    "pdf.fonttype": 42,
    "ps.fonttype": 42,
})


# =============================================================================
# GENERAL PLOTTING HELPERS
# =============================================================================

def style_axis(ax, grid=True):
    """
    Apply minimal consistent journal-style formatting.
    """

    ax.spines["left"].set_color(BLACK)
    ax.spines["bottom"].set_color(BLACK)

    ax.tick_params(
        direction="out",
        colors=BLACK
    )

    if grid:

        ax.grid(
            True,
            color=LIGHT_GREY,
            linewidth=0.45,
            alpha=0.32
        )

        ax.set_axisbelow(True)


def add_panel_title(ax, letter, title):
    """
    Fixed panel heading.

    Letter and title are deliberately separated.
    """

    ax.text(
        0.00,
        1.025,
        letter,
        transform=ax.transAxes,
        fontsize=12.5,
        fontweight="bold",
        ha="left",
        va="bottom",
        clip_on=False
    )

    ax.text(
        0.065,
        1.025,
        title,
        transform=ax.transAxes,
        fontsize=11.5,
        fontweight="normal",
        ha="left",
        va="bottom",
        clip_on=False
    )


def save_figure(fig, filename):
    """
    Save both raster and vector versions.
    """

    png = filename.with_suffix(".png")
    pdf = filename.with_suffix(".pdf")

    fig.savefig(
        png,
        dpi=600,
        bbox_inches="tight",
        pad_inches=0.05
    )

    fig.savefig(
        pdf,
        bbox_inches="tight",
        pad_inches=0.05
    )

    print(f"Saved: {png}")
    print(f"Saved: {pdf}")


# =============================================================================
# HAMMING BALL GEOMETRY
# =============================================================================

def calculate_hamming_ball_volumes():
    """
    V_q(L,r) = sum_{k=0}^r C(L,k)(q-1)^k
    """

    volumes = np.empty(
        L + 1,
        dtype=object
    )

    total = 0

    for r in range(L + 1):

        shell = (
            comb(L, r)
            * (q - 1) ** r
        )

        total += shell

        volumes[r] = total

    return volumes


HAMMING_VOLUMES = calculate_hamming_ball_volumes()

LOG10_HAMMING_VOLUMES = np.array([
    np.log10(float(v))
    for v in HAMMING_VOLUMES
])


# =============================================================================
# CHARACTERISTIC DISCOVERY RADIUS
# =============================================================================

def characteristic_radius_from_target_log10(log10_target):
    """
    Return smallest integer r satisfying

        log10 V_q(L,r) >= log10_target.

    If target <= 1, radius is zero.
    """

    if log10_target <= 0:
        return 0

    idx = np.searchsorted(
        LOG10_HAMMING_VOLUMES,
        log10_target,
        side="left"
    )

    return int(
        np.clip(
            idx,
            0,
            L
        )
    )


def characteristic_radius(log10_mu, log10_M=0):
    """
    Solve approximately

        V_q(L,r_c) ~ 1 / (mu M).

    Inputs are log10 values.
    """

    target = (
        -log10_mu
        - log10_M
    )

    return characteristic_radius_from_target_log10(
        target
    )


# =============================================================================
# RADIAL SUBSTITUTION DYNAMICS
# =============================================================================

def radial_drift(r):
    """
    E[Delta r | r]
        = 1 - q r / [L(q-1)]
        = 1 - r/r_eq
    """

    r = np.asarray(
        r,
        dtype=float
    )

    return 1 - r / R_EQ


def expected_radius(n):
    """
    Exact discrete-event expectation:

        E[R_n]
        = r_eq [1 - (1-alpha)^n]
    """

    n = np.asarray(
        n,
        dtype=float
    )

    return (
        R_EQ
        * (
            1
            - (1 - ALPHA) ** n
        )
    )


def mutation_events_for_radius(r):
    """
    Invert expected radial trajectory.
    """

    if r <= 0:
        return 0.0

    if r >= R_EQ:
        return np.inf

    return (
        np.log(1 - r / R_EQ)
        /
        np.log(1 - ALPHA)
    )


# =============================================================================
# MANUSCRIPT EXAMPLE VALUES
# =============================================================================

RC_SINGLE = characteristic_radius(
    log10_mu=-11,
    log10_M=0
)

RC_M1E4 = characteristic_radius(
    log10_mu=-11,
    log10_M=4
)

RC_M1E11 = characteristic_radius(
    log10_mu=-11,
    log10_M=11
)

NC_SINGLE = mutation_events_for_radius(
    RC_SINGLE
)

DRIFT_AT_RC = radial_drift(
    RC_SINGLE
)


# =============================================================================
# SANITY CHECKS
# =============================================================================

assert RC_SINGLE == 4
assert RC_M1E4 == 3
assert RC_M1E11 == 0

assert abs(R_EQ - 76) < 1e-12


# =============================================================================
# MONTE CARLO
# =============================================================================

def run_monte_carlo(
    n_trajectories=N_TRAJECTORIES,
    n_events=N_EVENTS,
    seed=RNG_SEED
):
    """
    Exact q-state substitution process.

    Each trajectory begins as the ancestral sequence:

        [0, 0, ..., 0]

    At every event:

        1. choose one of L sites uniformly;
        2. replace its current state by one of the other q-1 states
           uniformly.

    Hamming distance is measured from the ancestral all-zero sequence.
    """

    rng = np.random.default_rng(seed)

    sequences = np.zeros(
        (n_trajectories, L),
        dtype=np.uint8
    )

    distances = np.zeros(
        (n_events + 1, n_trajectories),
        dtype=np.uint8
    )

    current_distance = np.zeros(
        n_trajectories,
        dtype=np.int16
    )

    rows = np.arange(
        n_trajectories
    )

    for event in range(
        1,
        n_events + 1
    ):

        sites = rng.integers(
            0,
            L,
            size=n_trajectories
        )

        old = sequences[
            rows,
            sites
        ].copy()

        # Choose uniformly from q-1 states excluding current state.
        raw = rng.integers(
            0,
            q - 1,
            size=n_trajectories,
            dtype=np.uint8
        )

        new = (
            raw
            +
            (raw >= old).astype(np.uint8)
        )

        outward = (
            (old == 0)
            &
            (new != 0)
        )

        inward = (
            (old != 0)
            &
            (new == 0)
        )

        current_distance += (
            outward.astype(np.int16)
        )

        current_distance -= (
            inward.astype(np.int16)
        )

        sequences[
            rows,
            sites
        ] = new

        distances[event] = (
            current_distance.astype(np.uint8)
        )

    return distances


print()
print("Running Monte Carlo simulation...")

DISTANCES = run_monte_carlo()

print("Monte Carlo complete.")


EVENTS = np.arange(
    N_EVENTS + 1
)

ANALYTIC_MEAN = expected_radius(
    EVENTS
)

MC_MEAN = DISTANCES.mean(
    axis=1
)

MC_LOW = np.quantile(
    DISTANCES,
    LOW_Q,
    axis=1
)

MC_HIGH = np.quantile(
    DISTANCES,
    HIGH_Q,
    axis=1
)


# =============================================================================
# FIGURE 2A
# OUTWARD RADIAL DRIFT
# =============================================================================

def draw_figure_2a(ax):

    style_axis(
        ax,
        grid=True
    )

    r = np.linspace(
        0,
        80,
        1000
    )

    drift = radial_drift(r)

    # Main relationship
    ax.plot(
        r,
        drift,
        color=BLUE,
        linewidth=2.4,
        solid_capstyle="round"
    )

    # Zero drift
    ax.axhline(
        0,
        color=GREY,
        linewidth=0.8,
        linestyle="--"
    )

    # Equilibrium radius
    ax.axvline(
        R_EQ,
        color=GREY,
        linewidth=0.8,
        linestyle=":"
    )

    # Discovery-radius point
    ax.scatter(
        RC_SINGLE,
        DRIFT_AT_RC,
        s=38,
        color=ORANGE,
        edgecolor=WHITE,
        linewidth=0.7,
        zorder=5
    )

    # Equilibrium point
    ax.scatter(
        R_EQ,
        0,
        s=30,
        color=BLACK,
        zorder=5
    )

    # NO INTERNAL TEXT

    ax.set_xlim(
        0,
        80
    )

    ax.set_ylim(
        -0.07,
        1.05
    )

    ax.set_xlabel(
        "Hamming distance from ancestor, $r$"
    )

    ax.set_ylabel(
        r"Expected radial increment, "
        r"$\mathbb{E}[\Delta r\mid r]$"
    )

    ax.xaxis.set_major_locator(
        MultipleLocator(10)
    )

    ax.yaxis.set_major_locator(
        MultipleLocator(0.2)
    )

    add_panel_title(
        ax,
        "A",
        "Outward radial drift"
    )


# =============================================================================
# FIGURE 2B
# EXPECTED RADIAL TRAJECTORY
# =============================================================================

def draw_figure_2b(ax):

    style_axis(
        ax,
        grid=True
    )

    n = np.linspace(
        0,
        300,
        1500
    )

    r = expected_radius(n)

    ax.plot(
        n,
        r,
        color=BLUE,
        linewidth=2.4,
        solid_capstyle="round"
    )

    # Equilibrium guide only.
    ax.axhline(
        R_EQ,
        color=GREY,
        linewidth=0.8,
        linestyle=":"
    )

    # Manuscript discovery point.
    ax.scatter(
        NC_SINGLE,
        RC_SINGLE,
        s=38,
        color=ORANGE,
        edgecolor=WHITE,
        linewidth=0.7,
        zorder=5
    )

    # NO r_eq TEXT
    # NO n_c TEXT ON MAIN PANEL

    ax.set_xlim(
        0,
        300
    )

    ax.set_ylim(
        0,
        80
    )

    ax.set_xlabel(
        "Mutation events, $n$"
    )

    ax.set_ylabel(
        r"Expected Hamming distance, "
        r"$\mathbb{E}[R_n]$"
    )

    ax.xaxis.set_major_locator(
        MultipleLocator(50)
    )

    ax.yaxis.set_major_locator(
        MultipleLocator(10)
    )

    add_panel_title(
        ax,
        "B",
        "Expected radial trajectory"
    )

    # -------------------------------------------------------------------------
    # MINIMAL INSET
    # -------------------------------------------------------------------------

    inset = inset_axes(
        ax,
        width="31%",
        height="30%",
        loc="center right",
        borderpad=1.25
    )

    early_n = np.linspace(
        0,
        12,
        500
    )

    exact = expected_radius(
        early_n
    )

    # Exact trajectory
    inset.plot(
        early_n,
        exact,
        color=BLUE,
        linewidth=1.5
    )

    # Linear early-time approximation
    inset.plot(
        early_n,
        early_n,
        color=GREY,
        linewidth=1.0,
        linestyle="--"
    )

    # Discovery point
    inset.scatter(
        NC_SINGLE,
        RC_SINGLE,
        s=20,
        color=ORANGE,
        zorder=5
    )

    # Guides
    inset.plot(
        [NC_SINGLE, NC_SINGLE],
        [0, RC_SINGLE],
        color=ORANGE,
        linewidth=0.7,
        linestyle=":"
    )

    inset.plot(
        [0, NC_SINGLE],
        [RC_SINGLE, RC_SINGLE],
        color=ORANGE,
        linewidth=0.7,
        linestyle=":"
    )

    # NO TITLE
    # NO n_c LABEL
    # NO OTHER TEXT

    inset.set_xlim(
        0,
        12
    )

    inset.set_ylim(
        0,
        12
    )

    inset.set_xticks(
        [0, 4, 8, 12]
    )

    inset.set_yticks(
        [0, 4, 8, 12]
    )

    inset.set_xlabel(
        "$n$",
        fontsize=6.8,
        labelpad=0
    )

    inset.set_ylabel(
        "$r$",
        fontsize=6.8,
        labelpad=0
    )

    inset.tick_params(
        labelsize=6.3,
        length=2.5,
        width=0.6
    )

    inset.grid(
        True,
        color=LIGHT_GREY,
        linewidth=0.3,
        alpha=0.35
    )

    for spine in inset.spines.values():

        spine.set_visible(True)
        spine.set_linewidth(0.55)
        spine.set_color(GREY)


# =============================================================================
# FIGURE 2C
# MONTE CARLO
# =============================================================================

def draw_figure_2c(ax):

    style_axis(
        ax,
        grid=True
    )

    # Monte Carlo interval
    ax.fill_between(
        EVENTS,
        MC_LOW,
        MC_HIGH,
        color=PALE_BLUE,
        alpha=0.75,
        linewidth=0,
        label="Monte Carlo 5–95% interval"
    )

    # Monte Carlo mean
    ax.plot(
        EVENTS,
        MC_MEAN,
        color=ORANGE,
        linewidth=2.25,
        label="Monte Carlo mean"
    )

    # Analytic expectation
    ax.plot(
        EVENTS,
        ANALYTIC_MEAN,
        color=BLUE,
        linewidth=1.65,
        linestyle="--",
        label="Analytic expectation"
    )

    # Equilibrium guide
    ax.axhline(
        R_EQ,
        color=GREY,
        linewidth=0.8,
        linestyle=":"
    )

    # NO EQUILIBRIUM TEXT

    ax.set_xlim(
        0,
        300
    )

    ax.set_ylim(
        0,
        80
    )

    ax.set_xlabel(
        "Mutation events, $n$"
    )

    ax.set_ylabel(
        "Hamming distance from ancestor"
    )

    ax.xaxis.set_major_locator(
        MultipleLocator(50)
    )

    ax.yaxis.set_major_locator(
        MultipleLocator(10)
    )

    add_panel_title(
        ax,
        "C",
        "Monte Carlo validation"
    )

    # Simple legend in empty lower-right region
    ax.legend(
        loc="lower right",
        frameon=False,
        handlelength=2.8,
        borderaxespad=0.7
    )


# =============================================================================
# FIGURE 3A DATA
# =============================================================================

RARITY = np.linspace(
    0,
    60,
    2000
)

RC_BY_RARITY = np.array([
    characteristic_radius_from_target_log10(x)
    for x in RARITY
])


# =============================================================================
# FIGURE 3A
# FUNCTIONAL RARITY
# =============================================================================

def draw_figure_3a(ax):

    style_axis(
        ax,
        grid=True
    )

    ax.step(
        RARITY,
        RC_BY_RARITY,
        where="post",
        color=BLUE,
        linewidth=2.25
    )

    # Manuscript value mu = 1e-11
    ax.axvline(
        11,
        color=ORANGE,
        linewidth=1.0,
        linestyle=":"
    )

    ax.scatter(
        11,
        RC_SINGLE,
        s=42,
        color=ORANGE,
        edgecolor=WHITE,
        linewidth=0.7,
        zorder=5
    )

    # NO TEXT INSIDE PANEL

    ax.set_xlim(
        0,
        60
    )

    ax.set_ylim(
        0,
        32
    )

    ax.set_xlabel(
        r"Functional rarity, $-\log_{10}\mu$"
    )

    ax.set_ylabel(
        r"Characteristic radius, $r_c$"
    )

    ax.xaxis.set_major_locator(
        MultipleLocator(10)
    )

    ax.yaxis.set_major_locator(
        MultipleLocator(5)
    )

    add_panel_title(
        ax,
        "A",
        "Functional rarity sets the required radius"
    )


# =============================================================================
# FIGURE 3B DATA
# =============================================================================

RARITY_MAX = 32
LOGM_MAX = 13

x_heat = np.linspace(
    0,
    RARITY_MAX,
    800
)

y_heat = np.linspace(
    0,
    LOGM_MAX,
    500
)

X, Y = np.meshgrid(
    x_heat,
    y_heat
)

LOG_TARGET = X - Y

RADIUS_GRID = np.zeros_like(
    LOG_TARGET,
    dtype=int
)

positive = LOG_TARGET > 0

RADIUS_GRID[positive] = np.searchsorted(
    LOG10_HAMMING_VOLUMES,
    LOG_TARGET[positive],
    side="left"
)

RADIUS_GRID = np.clip(
    RADIUS_GRID,
    0,
    L
)


# =============================================================================
# FIGURE 3B
# PARALLELISM LANDSCAPE
# =============================================================================

def draw_figure_3b(ax):

    style_axis(
        ax,
        grid=False
    )

    max_radius = int(
        RADIUS_GRID.max()
    )

    # Discrete viridis palette
    base = plt.get_cmap(
        "viridis"
    )

    colours = base(
        np.linspace(
            0.12,
            0.93,
            max_radius + 1
        )
    )

    cmap = ListedColormap(
        colours
    )

    boundaries = np.arange(
        -0.5,
        max_radius + 1.5,
        1
    )

    norm = BoundaryNorm(
        boundaries,
        cmap.N
    )

    mesh = ax.pcolormesh(
        X,
        Y,
        RADIUS_GRID,
        cmap=cmap,
        norm=norm,
        shading="nearest",
        rasterized=True
    )

    # -------------------------------------------------------------------------
    # mu M = 1 boundary
    # -------------------------------------------------------------------------

    diagonal_max = min(
        RARITY_MAX,
        LOGM_MAX
    )

    diagonal = np.linspace(
        0,
        diagonal_max,
        300
    )

    ax.plot(
        diagonal,
        diagonal,
        color=WHITE,
        linewidth=1.2,
        linestyle="--",
        zorder=4
    )

    # -------------------------------------------------------------------------
    # Manuscript rarity slice: mu = 1e-11
    # -------------------------------------------------------------------------

    ax.axvline(
        11,
        color=WHITE,
        linewidth=0.85,
        linestyle=":",
        alpha=0.9,
        zorder=4
    )

    # -------------------------------------------------------------------------
    # Manuscript points
    #
    # NO LABELS.
    # -------------------------------------------------------------------------

    ax.scatter(
        11,
        0,
        s=48,
        color=BLUE,
        edgecolor=WHITE,
        linewidth=0.8,
        zorder=6,
        clip_on=False
    )

    ax.scatter(
        11,
        4,
        s=48,
        color=ORANGE,
        edgecolor=WHITE,
        linewidth=0.8,
        zorder=6
    )

    ax.scatter(
        11,
        11,
        s=48,
        color=GREEN,
        edgecolor=WHITE,
        linewidth=0.8,
        zorder=6
    )

    # NO TEXT INSIDE HEATMAP.
    # NO MU LABEL.
    # NO M LABELS.
    # NO R_C LABELS.
    # NO DIAGONAL LABEL.
    # NO CONTOUR LABELS.

    ax.set_xlim(
        0,
        RARITY_MAX
    )

    ax.set_ylim(
        0,
        LOGM_MAX
    )

    ax.set_xlabel(
        r"Functional rarity, $-\log_{10}\mu$"
    )

    ax.set_ylabel(
        r"Effective parallelism, $\log_{10}M$"
    )

    ax.xaxis.set_major_locator(
        MultipleLocator(5)
    )

    ax.yaxis.set_major_locator(
        MultipleLocator(2)
    )

    add_panel_title(
        ax,
        "B",
        "Parallelism reduces the required radius"
    )

    return mesh, max_radius


# =============================================================================
# INDIVIDUAL PANEL EXPORTS
# =============================================================================

print()
print("Generating individual panels...")


# -----------------------------------------------------------------------------
# Figure 2A
# -----------------------------------------------------------------------------

fig, ax = plt.subplots(
    figsize=(6.2, 4.6)
)

draw_figure_2a(ax)

fig.subplots_adjust(
    left=0.14,
    right=0.98,
    bottom=0.15,
    top=0.90
)

save_figure(
    fig,
    PANEL_DIR / "Figure_2A_v5"
)

plt.close(fig)


# -----------------------------------------------------------------------------
# Figure 2B
# -----------------------------------------------------------------------------

fig, ax = plt.subplots(
    figsize=(6.2, 4.6)
)

draw_figure_2b(ax)

fig.subplots_adjust(
    left=0.14,
    right=0.98,
    bottom=0.15,
    top=0.90
)

save_figure(
    fig,
    PANEL_DIR / "Figure_2B_v5"
)

plt.close(fig)


# -----------------------------------------------------------------------------
# Figure 2C
# -----------------------------------------------------------------------------

fig, ax = plt.subplots(
    figsize=(6.2, 4.6)
)

draw_figure_2c(ax)

fig.subplots_adjust(
    left=0.14,
    right=0.98,
    bottom=0.15,
    top=0.90
)

save_figure(
    fig,
    PANEL_DIR / "Figure_2C_v5"
)

plt.close(fig)


# -----------------------------------------------------------------------------
# Figure 3A
# -----------------------------------------------------------------------------

fig, ax = plt.subplots(
    figsize=(6.8, 4.8)
)

draw_figure_3a(ax)

fig.subplots_adjust(
    left=0.13,
    right=0.98,
    bottom=0.15,
    top=0.90
)

save_figure(
    fig,
    PANEL_DIR / "Figure_3A_v5"
)

plt.close(fig)


# -----------------------------------------------------------------------------
# Figure 3B
# -----------------------------------------------------------------------------

fig, ax = plt.subplots(
    figsize=(7.0, 4.8)
)

mesh, max_radius = draw_figure_3b(ax)

cbar = fig.colorbar(
    mesh,
    ax=ax,
    pad=0.025,
    fraction=0.045,
    ticks=np.arange(
        0,
        max_radius + 1,
        2
    )
)

cbar.set_label(
    r"Characteristic radius, $r_c$"
)

cbar.outline.set_linewidth(
    0.7
)

fig.subplots_adjust(
    left=0.13,
    right=0.90,
    bottom=0.15,
    top=0.90
)

save_figure(
    fig,
    PANEL_DIR / "Figure_3B_v5"
)

plt.close(fig)


# =============================================================================
# COMBINED FIGURE 2
# =============================================================================

print()
print("Generating combined Figure 2...")

fig, axes = plt.subplots(
    1,
    3,
    figsize=(17.4, 5.0)
)

draw_figure_2a(
    axes[0]
)

draw_figure_2b(
    axes[1]
)

draw_figure_2c(
    axes[2]
)

fig.subplots_adjust(
    left=0.055,
    right=0.995,
    bottom=0.16,
    top=0.90,
    wspace=0.26
)

save_figure(
    fig,
    ROOT
    / "Figure_2_radial_substitution_dynamics_v5"
)

plt.close(fig)


# =============================================================================
# COMBINED FIGURE 3
# =============================================================================

print()
print("Generating combined Figure 3...")

fig, axes = plt.subplots(
    1,
    2,
    figsize=(14.2, 5.0),
    gridspec_kw={
        "width_ratios": [
            1.0,
            1.03
        ]
    }
)

draw_figure_3a(
    axes[0]
)

mesh, max_radius = draw_figure_3b(
    axes[1]
)

cbar = fig.colorbar(
    mesh,
    ax=axes[1],
    pad=0.025,
    fraction=0.045,
    ticks=np.arange(
        0,
        max_radius + 1,
        2
    )
)

cbar.set_label(
    r"Characteristic radius, $r_c$"
)

cbar.outline.set_linewidth(
    0.7
)

fig.subplots_adjust(
    left=0.065,
    right=0.955,
    bottom=0.16,
    top=0.90,
    wspace=0.25
)

save_figure(
    fig,
    ROOT
    / "Figure_3_discovery_geometry_v5"
)

plt.close(fig)


# =============================================================================
# EXPORT NUMERICAL DATA
# =============================================================================

with (
    DATA_DIR
    / "hamming_ball_volumes.csv"
).open(
    "w",
    newline=""
) as f:

    writer = csv.writer(f)

    writer.writerow([
        "radius",
        "hamming_ball_volume",
        "log10_volume"
    ])

    for r, volume in enumerate(
        HAMMING_VOLUMES
    ):

        writer.writerow([
            r,
            str(volume),
            np.log10(float(volume))
        ])


with (
    DATA_DIR
    / "monte_carlo_summary.csv"
).open(
    "w",
    newline=""
) as f:

    writer = csv.writer(f)

    writer.writerow([
        "mutation_events",
        "analytic_expectation",
        "monte_carlo_mean",
        "monte_carlo_q05",
        "monte_carlo_q95"
    ])

    for i in range(
        len(EVENTS)
    ):

        writer.writerow([
            int(EVENTS[i]),
            float(ANALYTIC_MEAN[i]),
            float(MC_MEAN[i]),
            float(MC_LOW[i]),
            float(MC_HIGH[i])
        ])


# =============================================================================
# NUMERICAL SUMMARY
# =============================================================================

summary = f"""
Protein sequence-space figure calculations
==========================================

L = {L}
q = {q}

Functional density:
mu = {MU:.3e}

Nucleotide mutation rate:
mu_nt = {MU_NT:.3e}

Missense fraction:
f_missense = 392/549 = {F_MISSENSE:.8f}

Whole-protein amino-acid-changing mutation rate:
U = {U:.8e} per generation

Radial equilibrium:
r_eq = {R_EQ:.6f}

Characteristic radii at mu = 1e-11:

M = 1:
r_c = {RC_SINGLE}

M = 1e4:
r_c = {RC_M1E4}

M = 1e11:
r_c = {RC_M1E11}

Single-trajectory mutation-event threshold:
n_c = {NC_SINGLE:.8f}

Expected radial increment at r_c = 4:
E[Delta r | r_c] = {DRIFT_AT_RC:.8f}

Monte Carlo:
trajectories = {N_TRAJECTORIES}
events = {N_EVENTS}
seed = {RNG_SEED}
interval = {LOW_Q:.2f} to {HIGH_Q:.2f}
"""


with (
    DATA_DIR
    / "numerical_summary.txt"
).open(
    "w"
) as f:

    f.write(summary)


# =============================================================================
# TERMINAL SUMMARY
# =============================================================================

print()
print("=" * 72)
print("V5 COMPLETE")
print("=" * 72)

print()
print(f"Output folder:")
print(f"  {ROOT.resolve()}")

print()
print("Main figures:")
print(
    "  Figure_2_radial_substitution_dynamics_v5.png/pdf"
)
print(
    "  Figure_3_discovery_geometry_v5.png/pdf"
)

print()
print("Numerical checks:")
print(
    f"  r_eq = {R_EQ:.0f}"
)
print(
    f"  r_c(mu=1e-11, M=1) = {RC_SINGLE}"
)
print(
    f"  r_c(mu=1e-11, M=1e4) = {RC_M1E4}"
)
print(
    f"  r_c(mu=1e-11, M=1e11) = {RC_M1E11}"
)
print(
    f"  n_c = {NC_SINGLE:.4f}"
)
print(
    f"  E[Delta r | r_c] = {DRIFT_AT_RC:.4f}"
)

print()
print("Finished.")

# %% [code]
```
