## Supplementary figures and images for "Coverage Geometry and Heuristic Discovery Times in Protein Sequence Space"

### Figure_2_radial_substitution_dynamics_v5.pdf

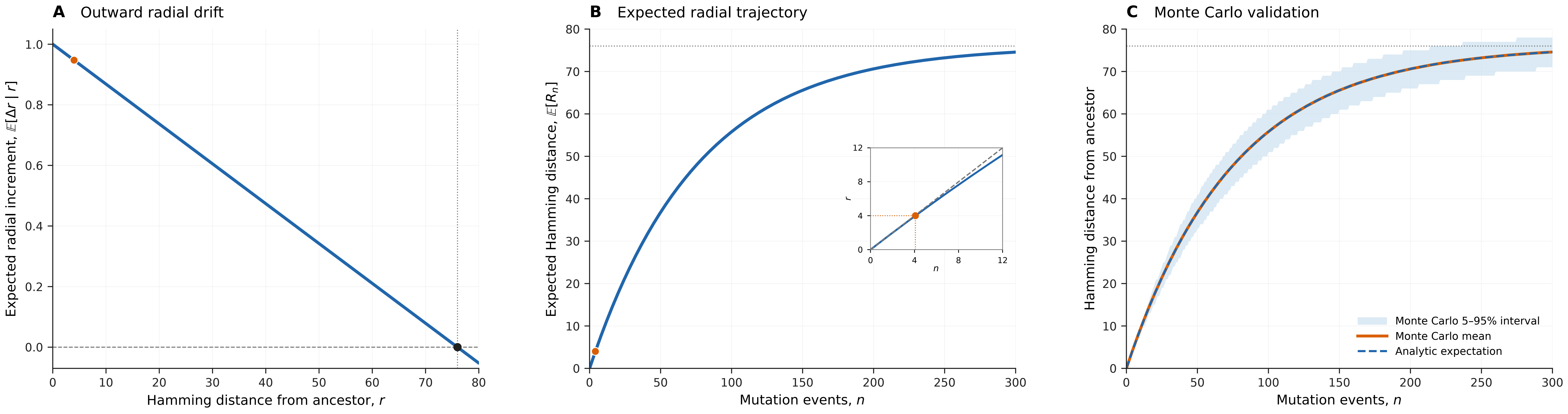

### Figure_2_radial_substitution_dynamics_v5.png

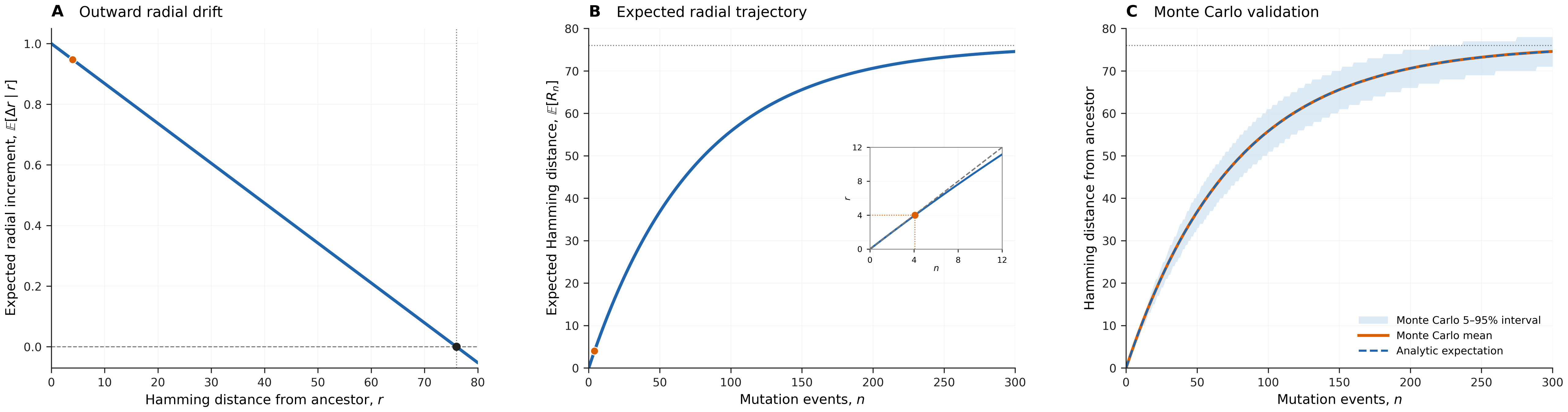

### Figure_2A_v5.pdf

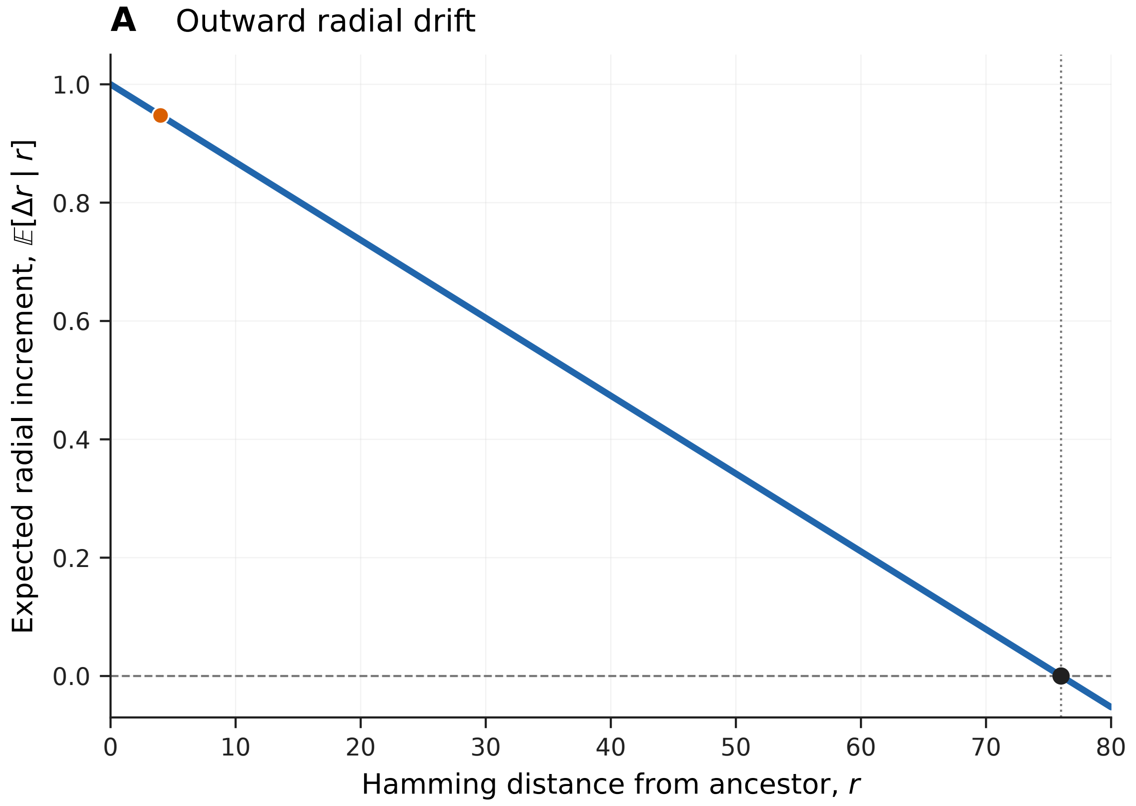

### Figure_2A_v5.png

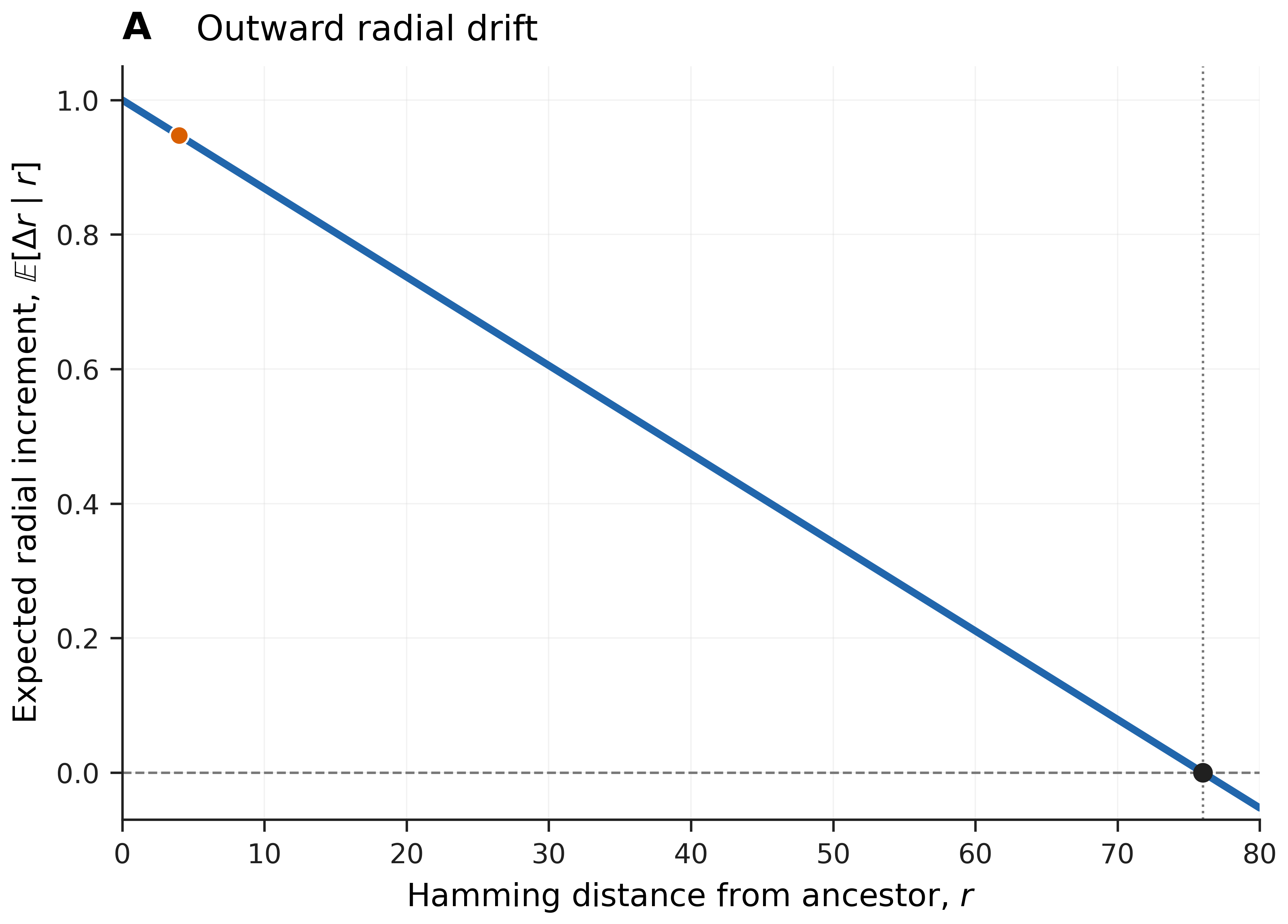

### Figure_2B_v5.pdf

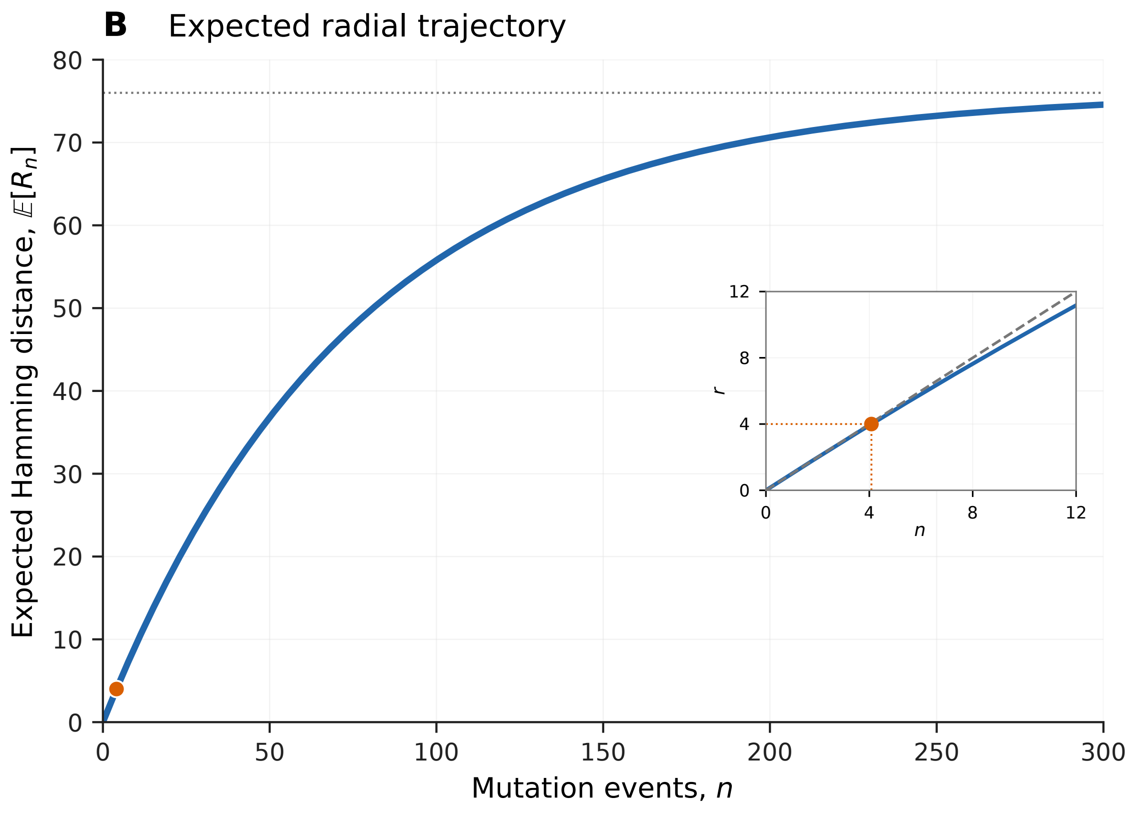

### Figure_2B_v5.png

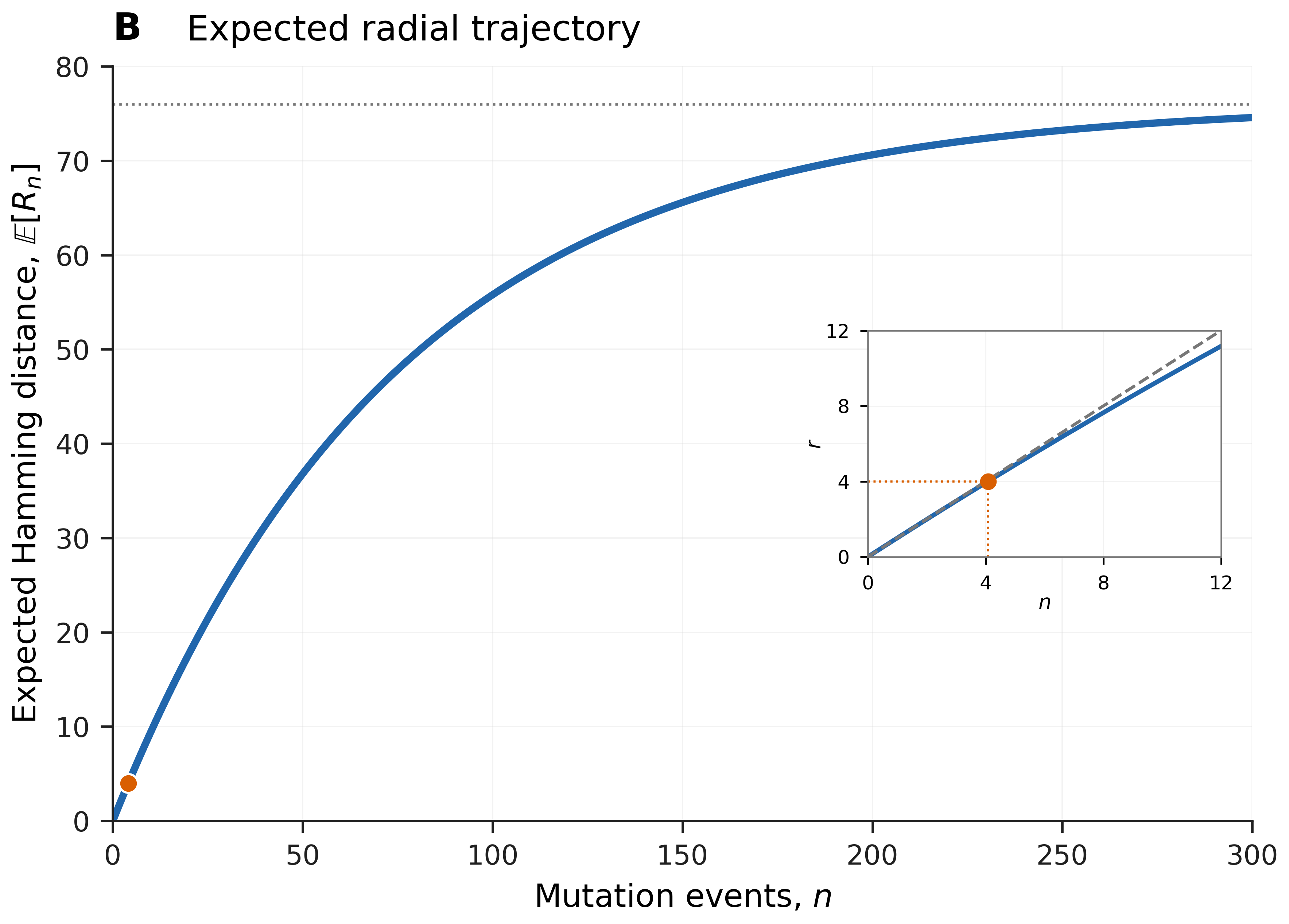

### Figure_2C_v5.pdf

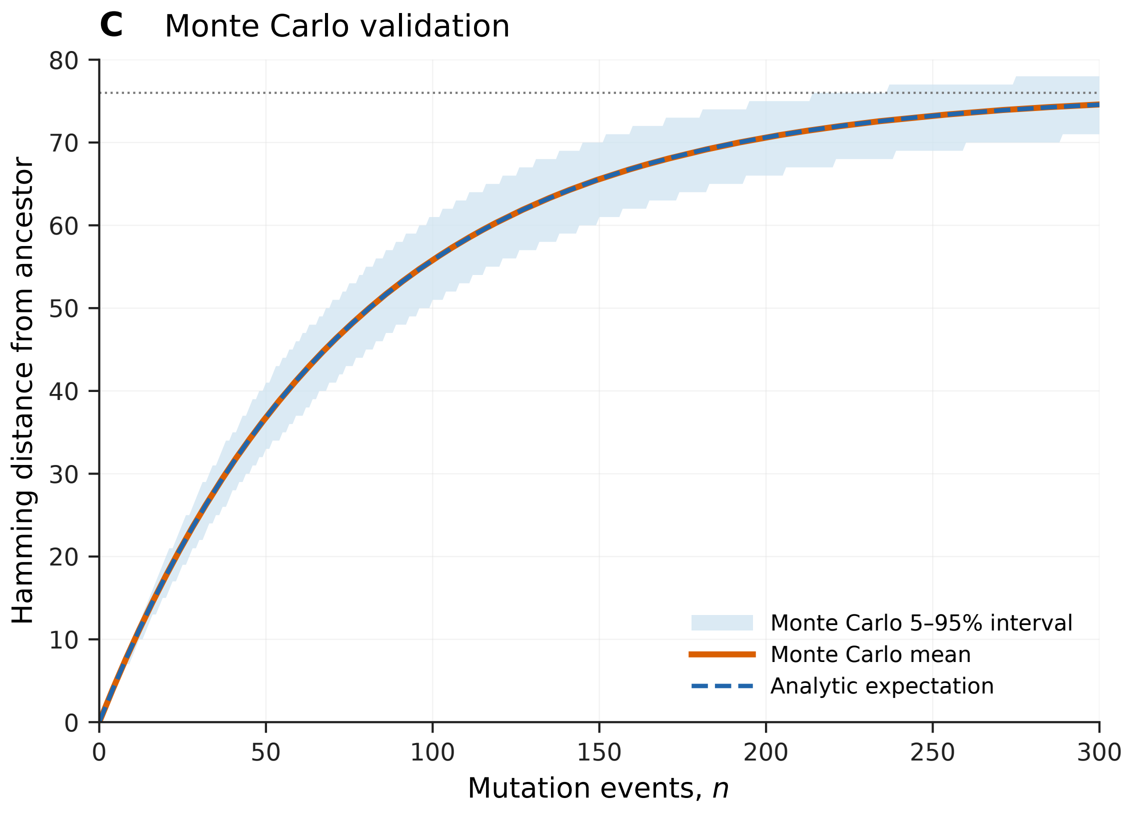

### Figure_2C_v5.png

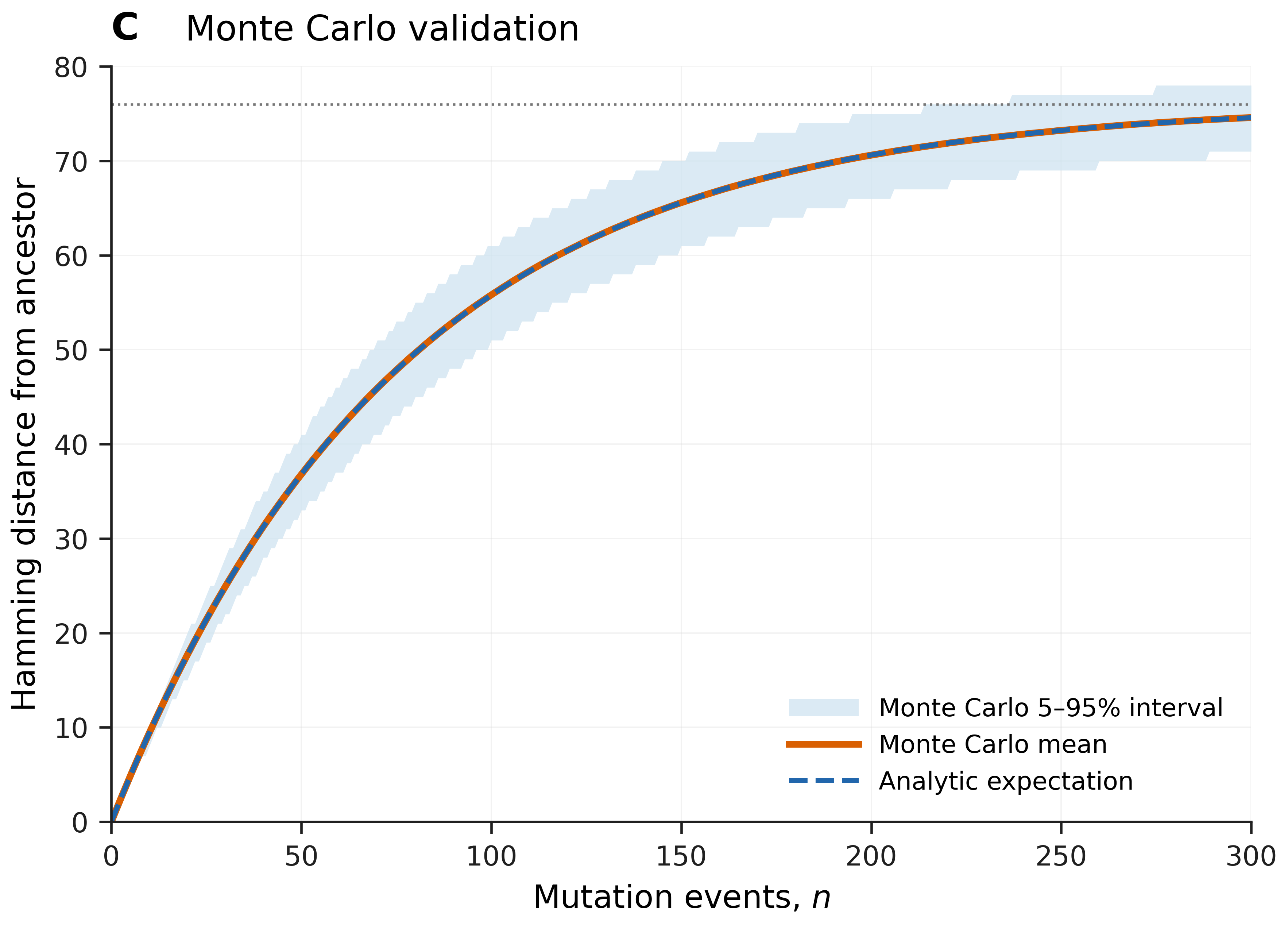

### Figure_3_discovery_geometry_v5.pdf

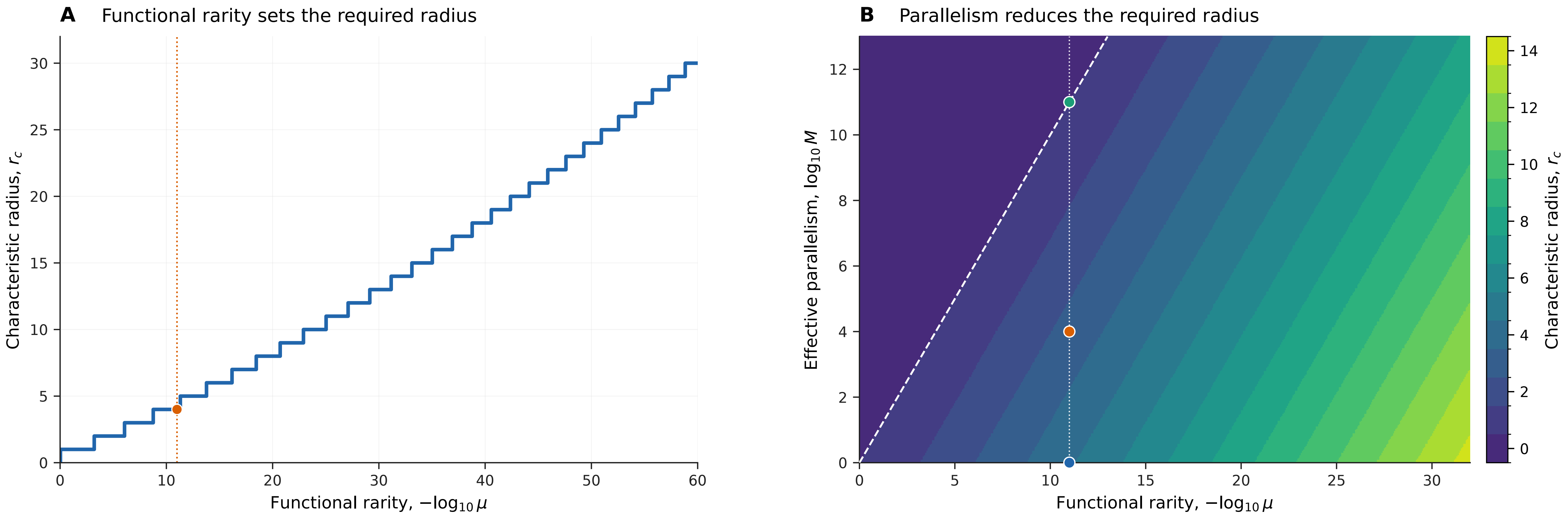

### Figure_3_discovery_geometry_v5.png

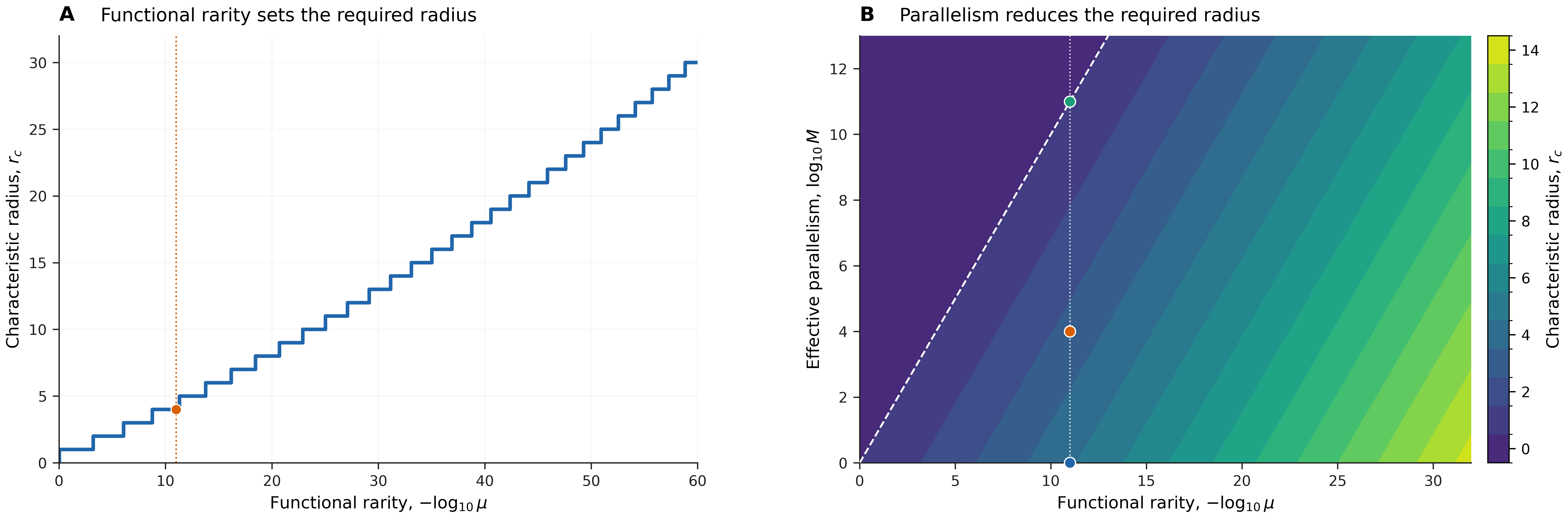

### Figure_3A_v5.pdf

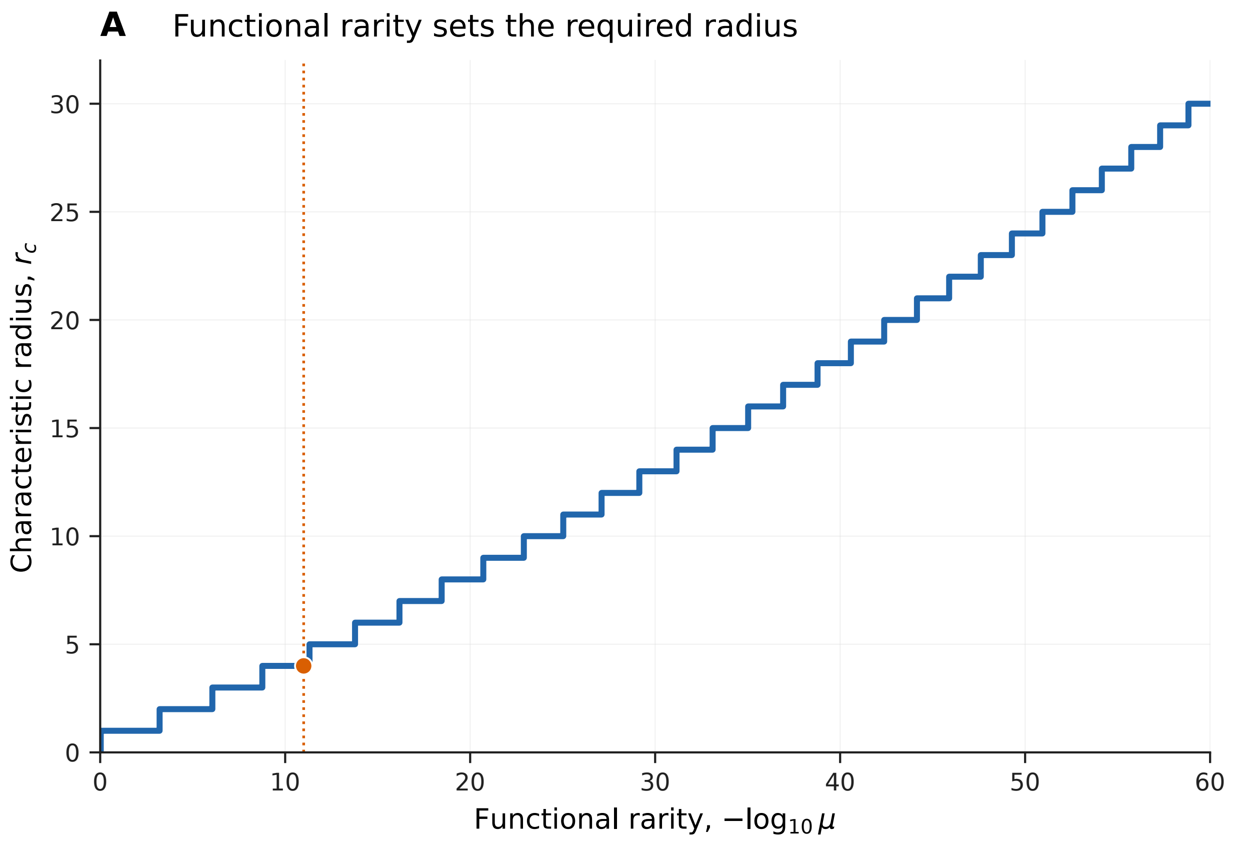

### Figure_3A_v5.png

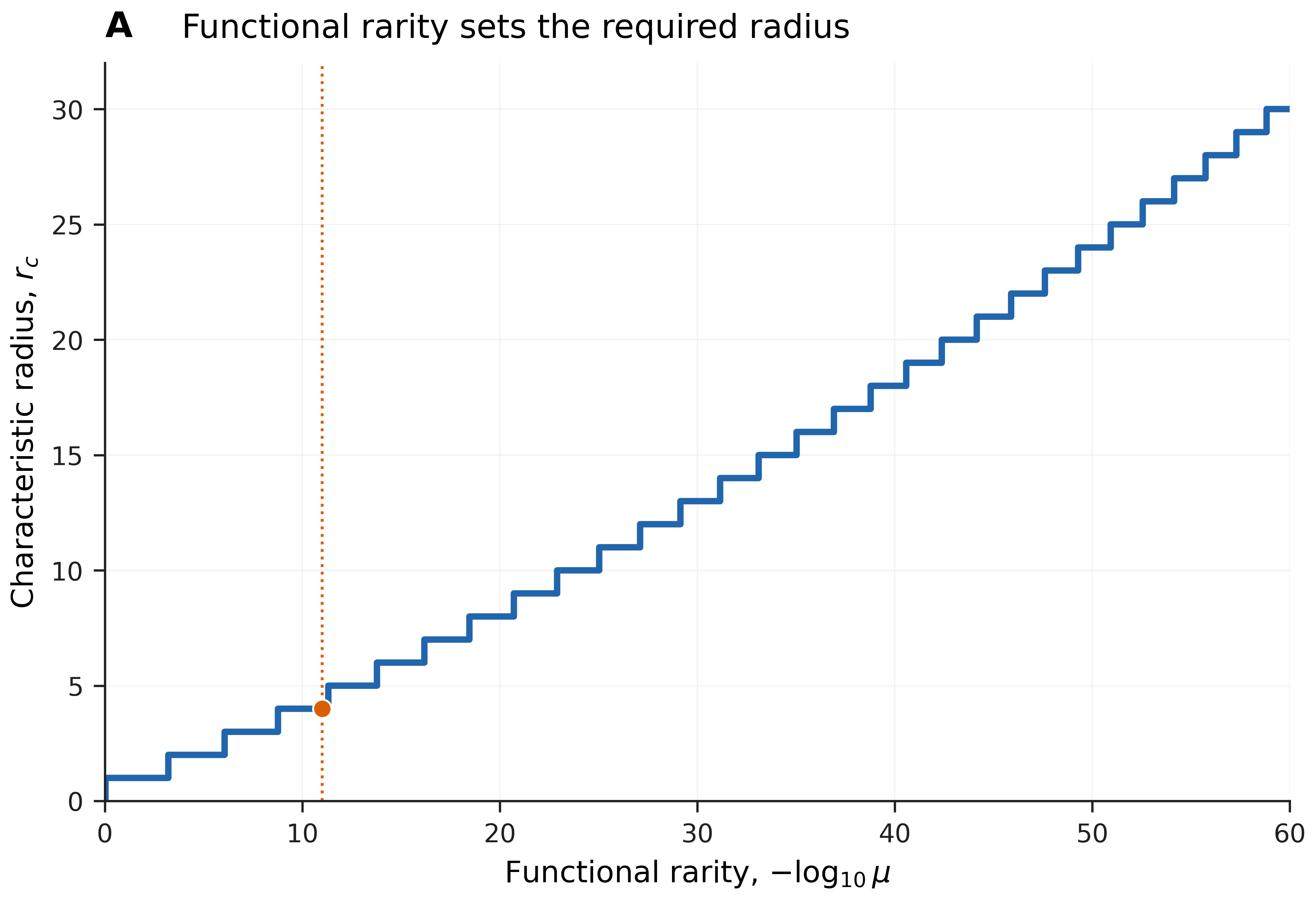

### Figure_3B_v5.pdf

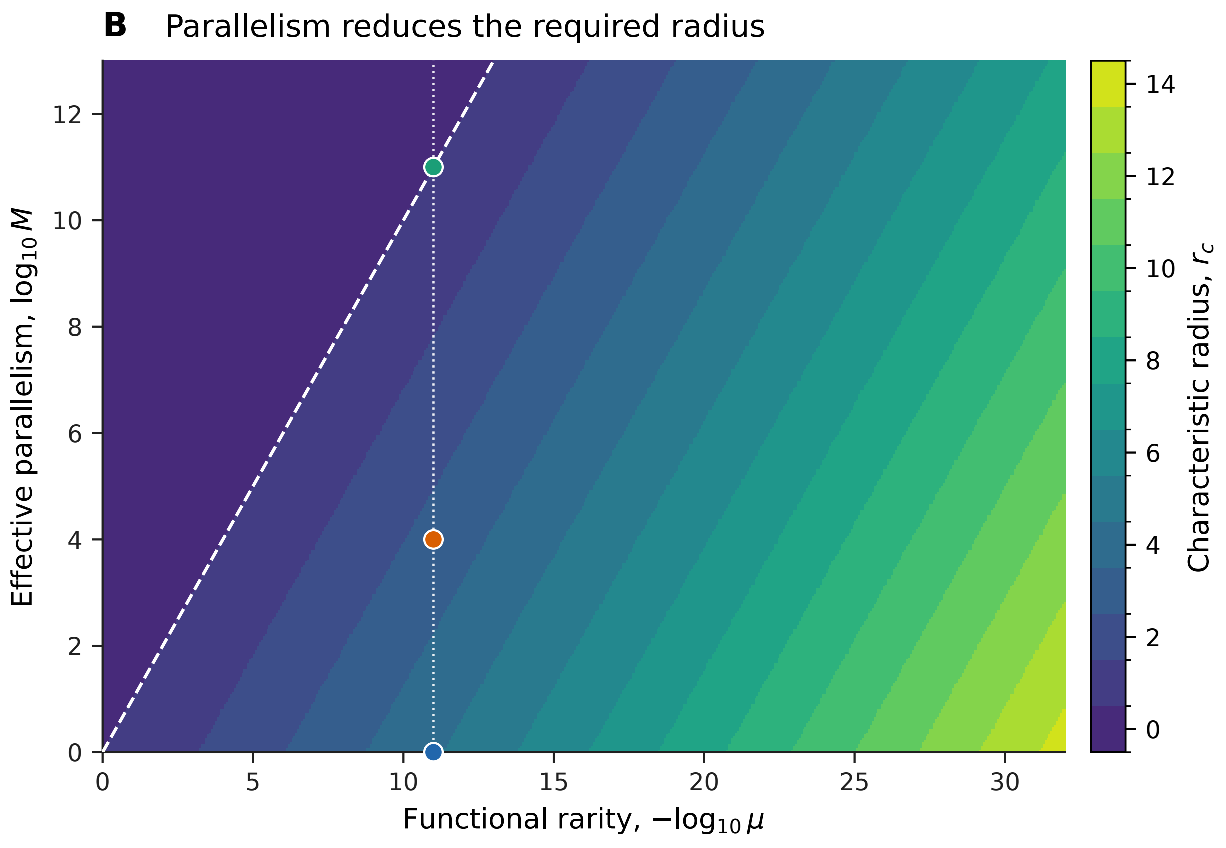

### Figure_3B_v5.png

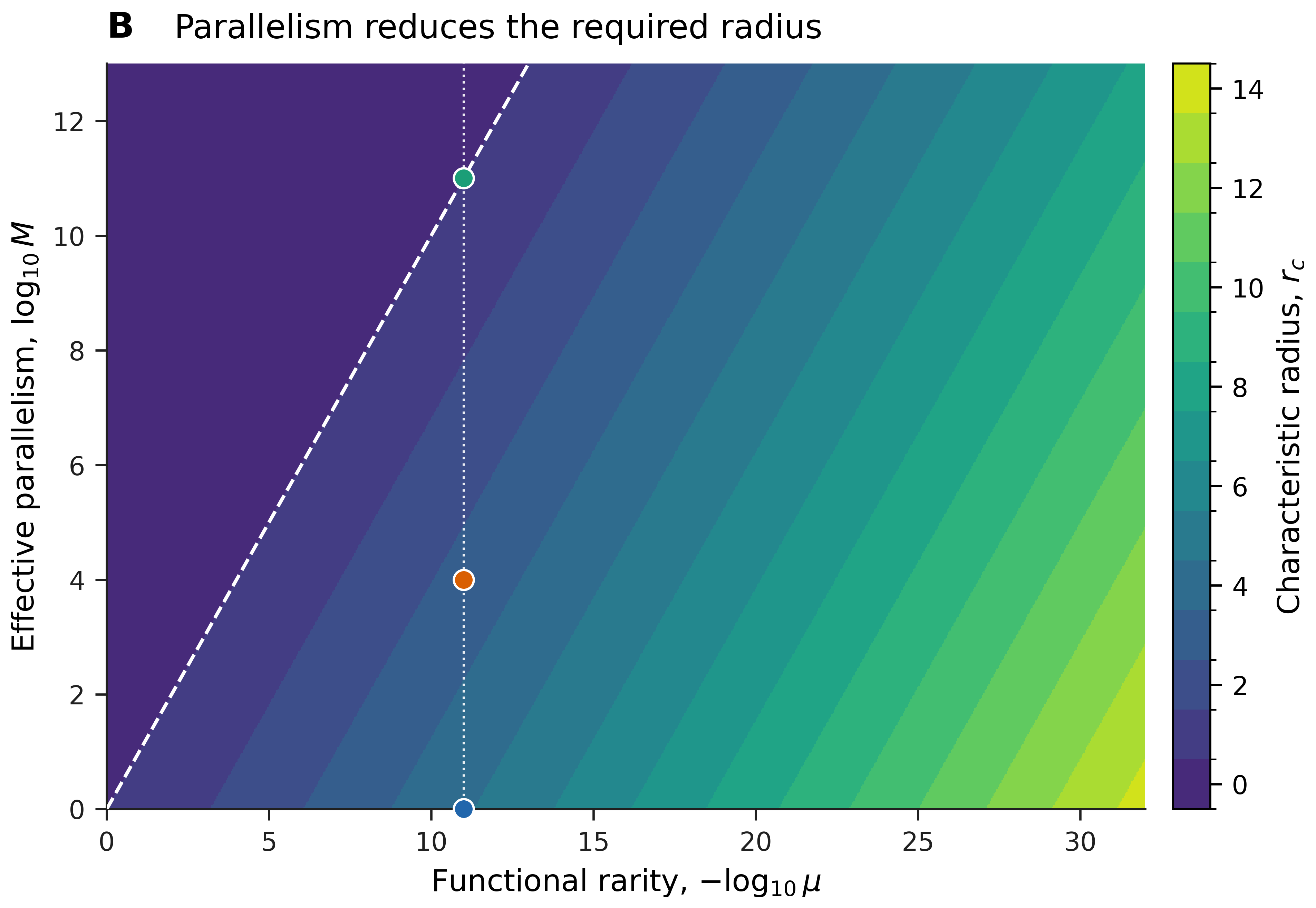
